# HSPD1 promotes neuroblastoma by augmenting MYCN expression

**DOI:** 10.64898/2026.09.03.749066

**Authors:** Shatha S Alassam, Dinesh Babu Manikandan, Hila Ben-David, Linor Cohen, Shai Kaluski-Kopatch, Laura Hruby, Shiran Dror, Poul Sorensen, Alberto Delaidelli, Gabriel Leprivier, Moshe Elkabets, Barak Rotblat

## Abstract

*MYCN* amplification is associated with poor outcomes in neuroblastoma (NB). *MYCN* encodes a transcription factor that induces the expression of mitochondrial one-carbon enzymes and chaperones, promoting metabolic reprogramming and aggressiveness in NB tumor cells. While the contribution of MYCN to changes in the mitochondrial proteome is well established, how mitochondrial proteostasis contributes to MYCN activity remains poorly understood. Mitochondrial HSP60 (HSPD1) is a conserved mitochondrial chaperone promoting the mitochondrial one-carbon pathway and mitochondrial translation by folding the one-carbon enzyme MTHFD2 and preventing aggregation of mitochondrial ribosomal proteins. Here, we show that HSPD1 depletion in *MYCN*-amplified NB cells led to reduced mitochondrial translation, *MYCN* expression, and tumorigenesis in vitro and in vivo. In addition, HSPD1 mRNA and protein correlate with *MYCN* expression in NB tumors. These results establish HSPD1 as a promising target for the treatment of *MYCN*-amplified NB.

## Introduction

Neuroblastoma (NB) is the most common extracranial solid tumor in children, with high-risk cases accounting for approximately 15% of all childhood cancer deaths. MYCN amplification occurs in up to 40% of high-risk cases and serves as a marker of poor prognosis, characterized by rapid disease progression, and an overall survival rate of less than 50% ^1^. MYCN encodes a transcription factor that promotes tumor development by increasing cell proliferation, inhibiting neuronal differentiation, and enhancing resistance to apoptosis ^2,3^. To sustain growth, MYCN induces metabolic reprogramming by upregulating the expression of nuclear-encoded mitochondrial enzymes ^3–5^. Mitochondria play a crucial role in NB, and mitochondrial inhibitors, particularly those targeting mitochondrial ribosomes, affect neuroblastoma differentiation and growth ^6–8^.

Most mitochondrial proteins are encoded in the nucleus, translated in the cytosol, and imported into the mitochondria as linear peptides. The mitochondrial HSP70 (mtHSP70/HSPA9) system first captures the translocating chain, then hands it off to the HSPD1/HSPE1 (HSP60/HSP10) chaperonin complex^9,10^. HSPD1 acts as a nanocage that encapsulates the unfolded substrate, providing an isolated environment in which it can fold ^11–13^. In cases where the demand for folding exceeds the chaperoning capacity of the mitochondria, the transcription factor ATF5 activates the mitochondrial unfolded protein response (mtUPR) ^14^, which includes the upregulation of HSPD1 expression^15^, to resolve the proteostasis crisis in the mitochondria^16^. HSPD1 is not only induced by mtUPR but is also essential for mounting the mtUPR, which is important for supporting tumorigenesis in prostate cancer cells^17^. Importantly, ATF5 was found to promote NB ^18^, and *HSPD1* mRNA correlated with poor patient outcomes in NB ^6^. Nevertheless, the functional role and mechanism of action of HSPD1 in NB are poorly understood.

By increasing the expression of mitochondrial proteins ^19^, MYCN is expected to increase the burden on the mitochondrial proteome quality control. It has been demonstrated that, in addition to mitochondrial enzymes, MYCN directly upregulates HSPD1 ^20^. On the other hand, inhibiting mitochondrial translation with doxycycline reduced the expression of several oncogenes, including c-MYC, in neuroblastoma ^8^. While it is known that MYCN alters mitochondrial metabolism in NB, the potential for retrograde signaling, in which mitochondrial proteostasis influences MYCN expression and activity, remains largely unexplored.

Here, we show that HSPD1 depletion reduces mitochondrial translation, MYCN expression, and tumorigenicity in *MYCN*-amplified neuroblastoma cells.

## Results

### HSPD1 is coexpressed with MYCN in NB tumors

HSPD1 is a MYCN transcriptional target gene ^20^ whose high expression correlates with poor patient outcomes in NB ^21^. To test whether *HSPD1* expression correlates with *MYCN* amplification status in patient tissue, we used the R2 platform and interrogated patient cohorts for which mRNA expression and MYCN status are available. We found significantly increased *HSPD1* mRNA expression in *MYCN*-amplified samples vs. non-amplified samples in three independent cohorts (Fig. 1a). In accord, we found a significant positive correlation between the levels of *HSPD1* and *MYCN* mRNA in four independent cohorts (Fig. 1b).

**Fig. 1.**
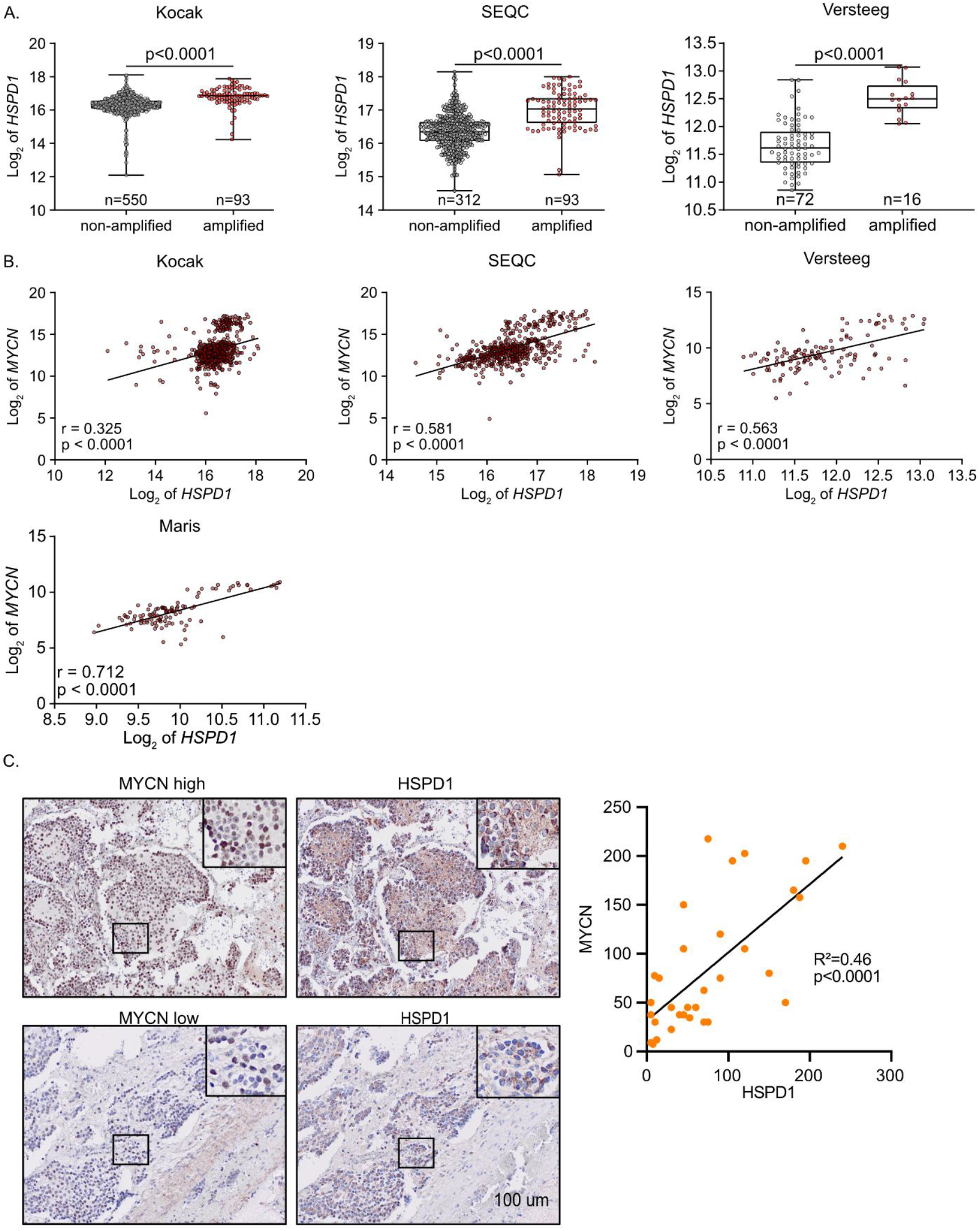
HSPD1 is correlated with MYCN in NB tumors. A. *HSPD1* expression in *MYCN*-amplified and nonamplified NB samples in the indicated patient-derived tumor RNAseq data was obtained from the indicated cohorts and analysed using R2 ^22^. B. HSPD1 and MYCN expression in the indicated patient-derived tumor RNAseq data was obtained from the indicated cohorts and analysed using R2 . C. HSPD1 and MYCN protein levels in NB tumor tissue were measured using IHC and anti-HSPD1 or MYCN antibodies. Staining was scored, and R and p were calculated using Spearman.

Using NB clinical samples, we found by IHC that HSPD1 is also coexpressed with MYCN at the protein level (Fig. 1c). Taken together, these data establish a significant correlation between HSPD1 and MYCN expression at the mRNA and protein levels in NB.

### HSPD1 promotes NB tumorigenicity in culture and in vivo

To test whether HSPD1 is functional in *MYCN*-amplified NB cells, KELLY and IMR-5, we attempted to generate an HSPD1 knockout (KO) using CRISPR-Cas9 system. However, isolated K clones did not grow into cell colonies, perhaps wich is likely attributed to the essential role of HSPD1 ^23^. We therefore selected a colony with low HSPD1 expression and used two shRNA targeting HSPD1 and lentiviruses to generate HSPD1 knockdown (KD) cells, as we did in ^24^.

We measured viability and colony formation (foci) in HSPD1 KD and control cells and found that HSPD1 KD reduced both parameters, indicating that HSPD1 promotes pro-tumorigenic traits in NB cell lines in culture (Fig. 2b and c).

**Fig. 2.**
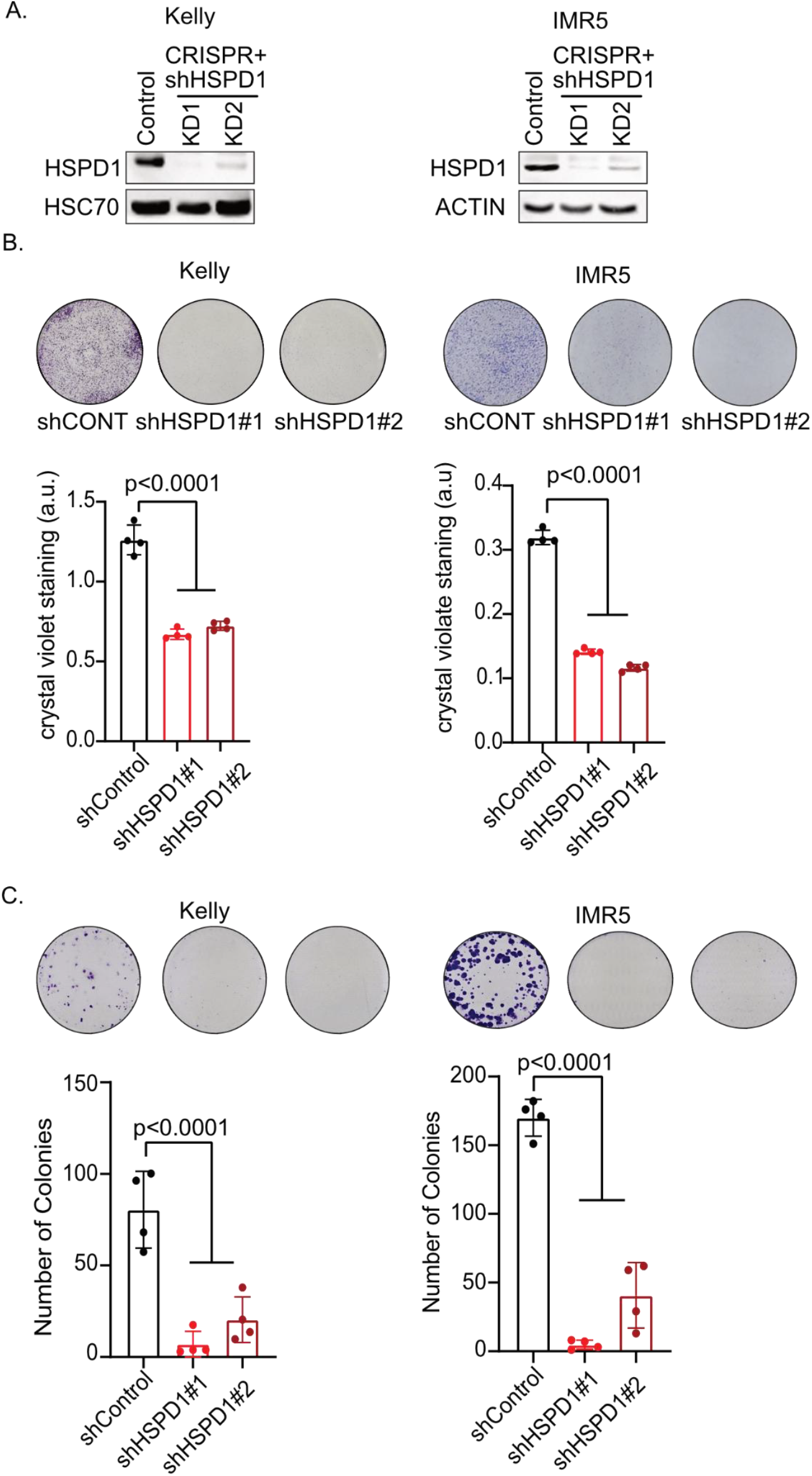
HSPD1 is functional in NB cells. A. HSPD1 expression in HSPD1 KD and control cells was tested using anti-HSPD1 antibodies. Actin or HSC70 was used as a loading control. B. HSPD1 KD and control cells were plated at equal numbers and grown for 5 days. Cells were stained with crystal violet, and plates were imaged. Typical images are shown. Crystal violet was dissolved in 10% acetic acid, and OD was measured. Bar graph represents the mean of independent biological experiments, error bars SD; n=4. C. HSPD1 KD and control cells were plated at low density and equal numbers and grown for 3 weeks. Cells were stained with crystal violet, and plates were imaged. Typical images are shown. Colony numbers were counted using ImageJ. Bar graph represents mean, error bars SD; n=4.

We tested HSPD1 function *in vivo* using xenograft and NOD-SCID gamma mice. We injected 8 million HSPD1 KD or control KELLY cells into the flanks of the mice and monitored tumor growth over time. We found delayed tumor growth in HSPD1 KD cells compared with controls (Fig. 3a), indicating HSPD1 KD reduced the tumorigenicity of NB cells in vivo.

**Fig. 3.**
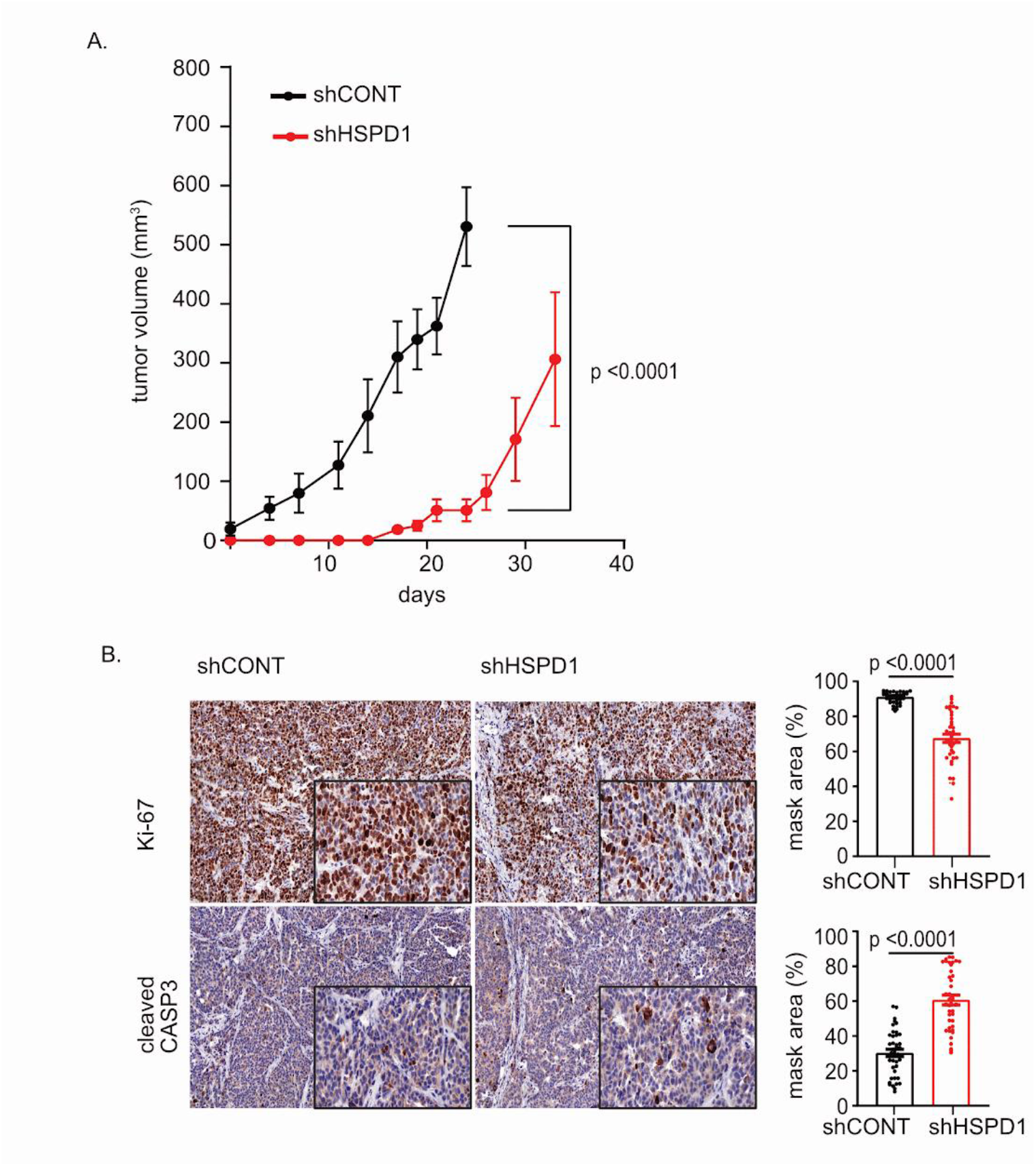
HSPD1 promotes NB tumor growth in mice. A. Eight million HSPD1 KD and control KELLY cells were injected into the flanks of NOD-SCID gamma mice. Tumors were measured using a caliper. n=5; unpaired t-test. B. Tumors were sectioned, stained with H&E and anti-Ki-67 or cleaved caspase-3 antibodies. Slides were scanned and staining quantified. Bar shows mean ± SD; Student’s T-test.

We measured Ki-67 and cleaved caspase-3, markers of proliferation and apoptosis respectivly, in HSPD1 KD and control tumors using IHC and found that HSPD1 KD reduced proliferation and increased apoptosis (Fig. 3b). These results indicate that HSPD1 promotes tumorigenicity in *MYCN*-amplified NB cells in vivo.

### Reduced *MYCN* mRNA and MYCN target genes in HSPD1 KD cells

To find how HSPD1 is linked to NB tumorigenicity, we profiled the transcriptomes of HSPD1 KD and control KELLY cells using RNAseq. Confirming KD efficiency, we found reduced *HSPD1* expression in HSPD1 KD cells (Fig. 4a). Notably, along with HSPD1 KD we found reduced *MYCN* expression (Fig. 4a). We found induction of *TP53*, and *MDM2*, which were also upregulated in HSPD1 KD A549 cells ^25^. The p53 targets *MIR34A* and *p21* were also upregulated in HSPD1 KD KELLY cells and previously shown in A549 cells ^25^. Consistent with earlier observations in A549 cells, U251 cells ^25^, and patient fibroblasts harboring HSPD1 mutations ^26^, HSPD1 knockdown in KELLY cells failed to activate the mitochondrial unfolded protein response (mtUPR), and ATF5, a marker of mtUPR ^27^, was downregulated in HSPD1 KD cells (Fig. 4a).

**Fig. 4.**
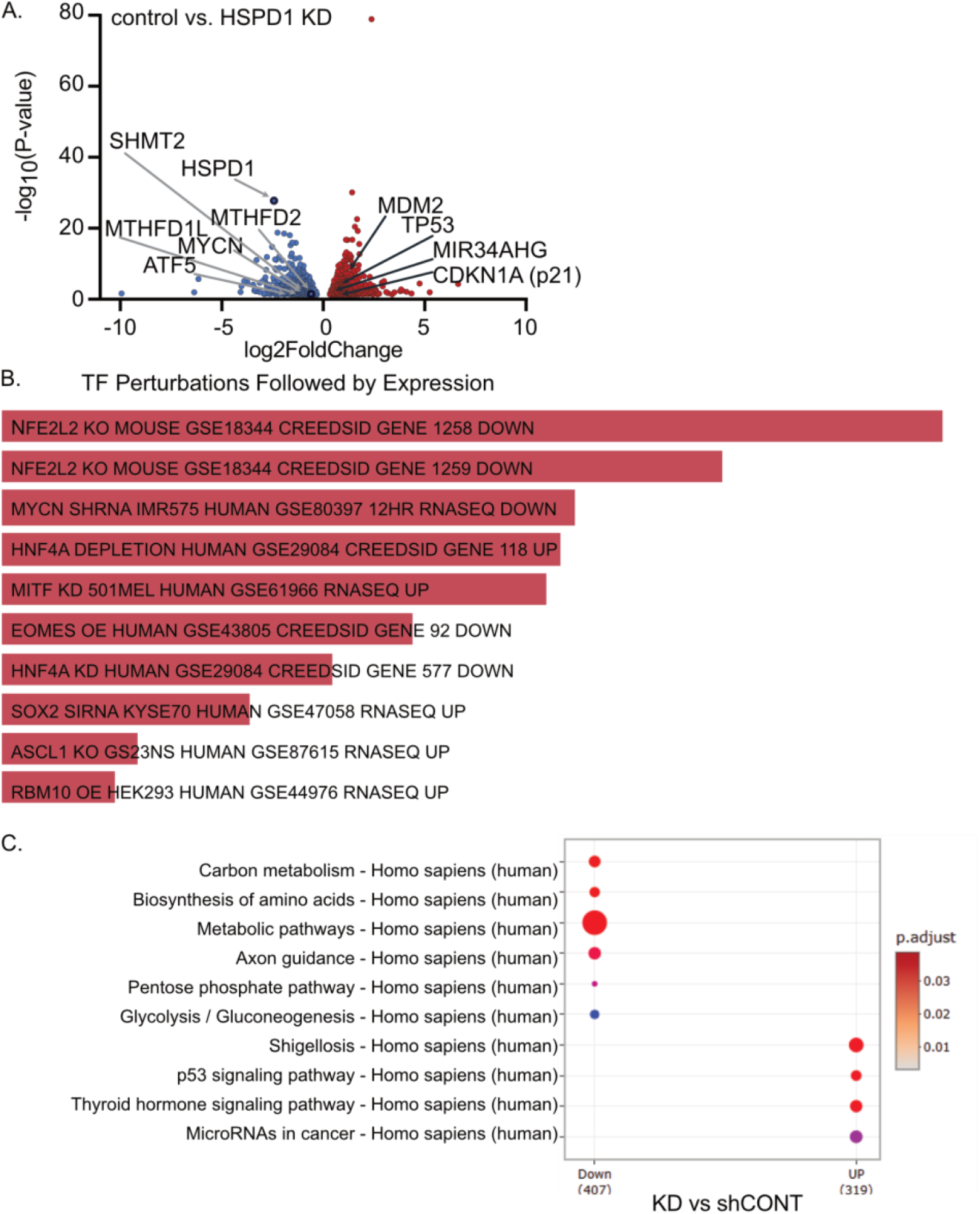
Transcriptome of control vs. HSPD1 KD in KELLY cells. A. RNAseq performed on HSPD1 KD and control KELLY cells (samples: 4 shCONT; 4 shHSPD1#1, 4 HSPD1#2). HSPD1 KD data were combined (n=8) and compared to shCONT. The volcano plot shows that differential genes and transcripts of interest are indicated. B. Differentially expressed gene lists were analyzed using ENRICHR. Genes affected by HSPD1 KD are similar to those found in MYCN KD experiments. C. KEGG analysis of genes whose expression was affected by HSPD1 KD vs. controls in KELLY cells.

To analyze the transcriptional effects of HSPD1 knockdown and identify differentially expressed genes, we preformed Enrichr web-based enrichment analysis ^28–30^. Within the category Transcription Factor Perturbations Followed by Expression, the analysis revealed that these genes are significantly upregulated and overlap with those identified in an independent study where MYCN was knocked down in NB cells (Fig. 4b). MYCN upregulates one-carbon pathway enzymes ^5,19,31^ and we found downregulation of the MYCN targets *MTHFD2, MTHFD2L* and *SHMT2* in HSPD1 KD KELLY cells, aligning with reduced MYCN activity (Fig. 4a). This finding suggests that the transcriptional response to HSPD1 depletion resembles that of MYCN depletion, supporting the notion that HSPD1 plays an important role in promoting MYCN expression and that down-regulation of MYCN is a major biological outcome of HSPD1 depletion. In addition, HSPD1 KD resembled KD of the antioxidant-stress-response transcription factor NFE2L2 (NRF2) (Fig. 4b). These data are consistent with findings in HSPD1 KD A549 cells ^25^.

KEGG pathway enrichment analysis further supported these observations (Fig. 4c). Downregulated pathways included glycolysis, carbon metabolism, and amino acid biosynthesis. These are consistent with decreased metabolic and mitochondrial activity following HSPD1 KD ^25^. Conversely, upregulated pathways were linked to p53 signaling, thyroid hormone signaling, and microRNAs in cancer, consistent with the activation of stress-response signaling pathways.

### HSPD1 promotes MYCN in NB cells

Having found that HSPD1 KD reduces *MYCN* mRNA levels, we next asked whether this is also true at the protein level. Using Western blot and immunofluorescence (IF), we found that MYCN expression was significantly reduced in HSPD1 KD NB cells. These findings suggest that HSPD1 promotes MYCN expression in NB cells (Fig. 5a and b).

**Fig. 5.**
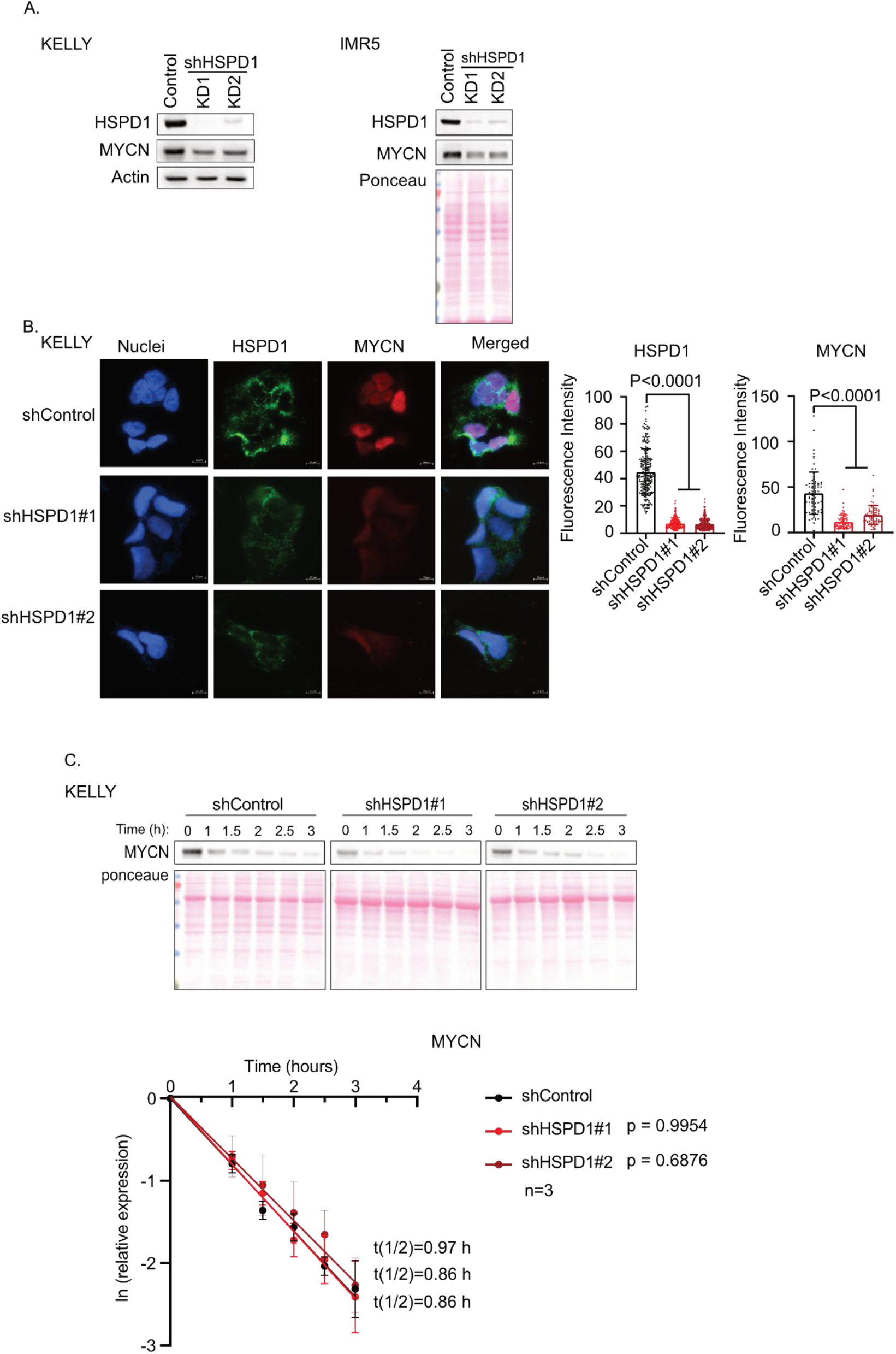
Reduced MYCN expression in HSPD1 KD NB cells. A. The indicated proteins in HSPD1 KD and control cells were analyzed by western blot. B. Control and HSPD1 KD cells were fixed and immunostained with anti-HSPD1 or anti-MYCN antibodies, with nuclei counterstained using DAPI. Images were acquired by fluorescence microscopy. Nuclear MYCN fluorescence intensity within DAPI-defined regions was quantified using ImageJ. Statistical significance was calculated using one-way ANOVA. C. HSPD1 KD and control cells were pulsed with CHX, and samples were collected at the indicated time points. MYCN levels were measured using Western blot and normalized to ponceau staining used as a loading control. Protein half-life was calculated using ln(normalized expression). P calculated using one-way ANOVA.

MYCN is regulated at the protein level by the ubiquitin-proteasome system ^32^. To test whether HSPD1 KD leads to reduced MYCN expression due to increased degradation, we used a cycloheximide (CHX) chase to measure the half-life of MYCN in control and HSPD1 KD KELLY cells. We found that HSPD1 KD did not significantly decrease the half-life of MYCN (Fig. 5c). These results indicate that HSPD1 does not affect MYCN half-life but does play a crucial role in promoting *MYCN* mRNA and protein expression.

Having found that HSPD1 KD KELLY cells exhibit delayed tumor growth in vivo (Fig. 3a), we wondered if the growing tumors showed reduced MYCN expression while maintaining HSPD1 KD. We used IHC to measure HSPD1 and MYCN levels in HSPD1 KD and control tumors and found that HSPD1 KD tumors exhibited reduced HSPD1 expression (Fig. S1). Surprisingly, we found moderate increased MYCN expression in HSPD1 KD tumors compared with controls (Fig. S1). These results suggest that in vivo, KELLY cells adapt to HSPD1 KD by upregulating MYCN expression, despite the effect on tumor growth.

### HSPD1 promotes mitochondrial translation in *MYCN*-amplified NB cells

To investigate a possible mechanism by which HSPD1 regulates MYCN expression, we turned our attention to mitochondrial translation. This is because MYCN expression was shown to depend on mitochondrial translation, and treating *MYCN*-amplified NB cells with mitochondrial translation inhibitors reduced MYCN expression ^8^. In addition, HSPD1 KD was shown to reduce mitochondrial translation and promote the aggregation of mitochondrial ribosomal proteins in HEK293 and yeast cells ^33^.

To measure mitochondrial translation in KELLY and IMR-5 *MYCN*-amplified NB cells, we pulsed HSPD1 KD and control cells with the synthetic clickable methionine analog L-homopropargylglycine (HPG) in the presence of cytosolic translation inhibitor emetine and labeled mitochondria using Mitotracker RED. We measured labeled HPG in the mitochondria and found reduced mitochondrial translation in HSPD1 KD cells (Fig. 6a and b). To test the specificity of labeling, we imaged cells without Emetine, without HPG, or with Emetine and chloramphenicol to inhibit both cytosolic and mitochondrial translation (Fig. S2). Our data show that HSPD1 promotes mitochondrial translation in *MYCN*-amplified NB cells.

**Fig. 6.**
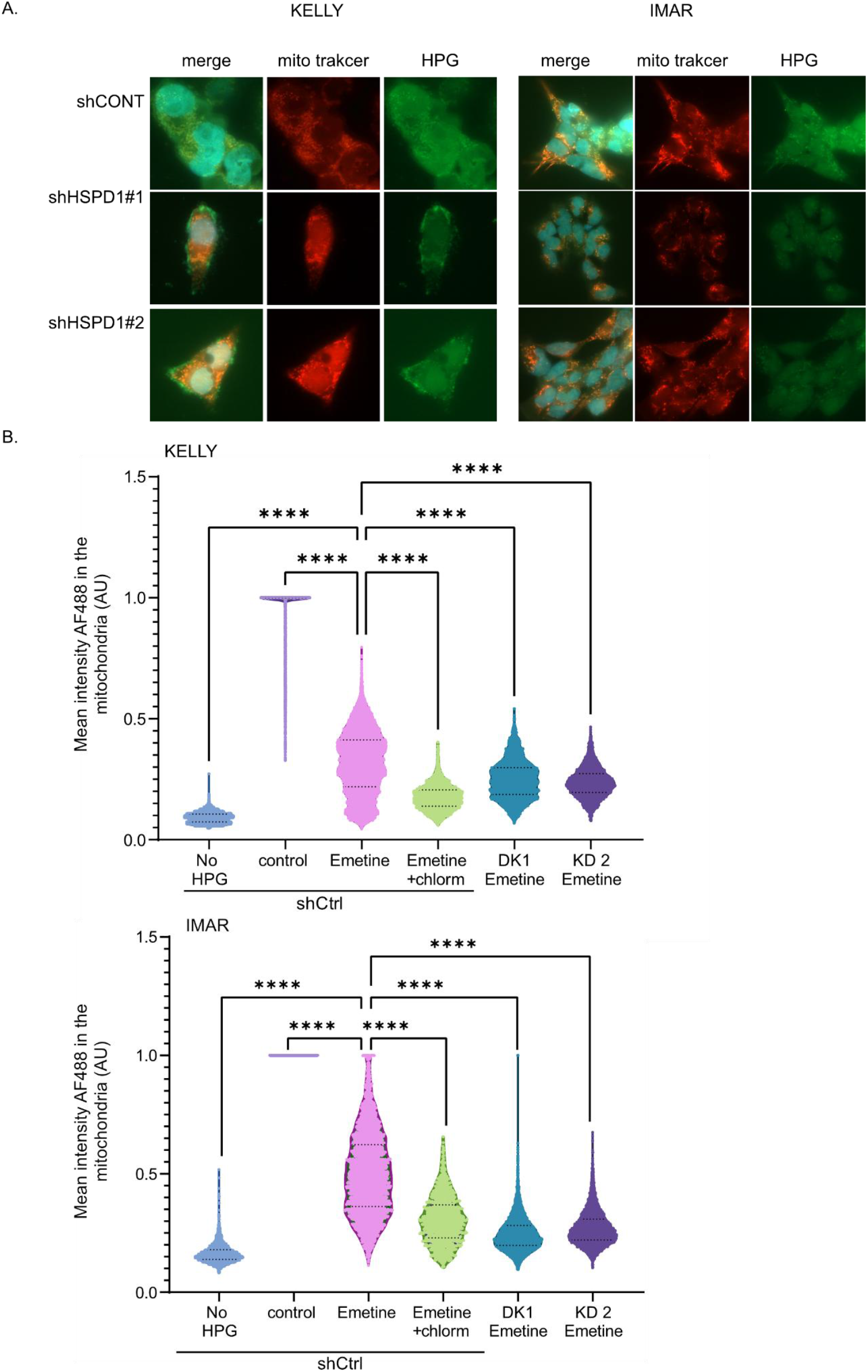
Reduced mitochondrial translation in HSPD1 KD NB cells. A. Representative fluorescence microscopy images of KELLY/ IMAR cells expressing control shRNA (shCtrl) or HSPD1 knockdown (KD1, KD2) treated with emetine (50 µg/mL) to inhibit cytosolic translation and isolate mitochondrial protein synthesis. Newly synthesized mitochondrial proteins were labeled with L-homopropargylglycine (HPG; green), mitochondria were counterstained with MitoTracker Red (red), and nuclei were stained with DAPI (blue). Scale bar = 10 µm. B. Quantification of mean HPG (Alexa Fluor 488) fluorescence intensity localized within mitochondrial regions across experimental conditions (in Arbitrary Units, A.U.). Data are presented as violin plots displaying the median (solid line) and quartiles (dashed lines). Statistical significance was calculated using a one-way ANOVA followed by Tukey’s [or Dunnett’s] multiple comparisons test (**** *p* < 0.0001).

## Discussion

The existence of a MYCN-mitochondria axis is well known in NB, as MYCN directly induced the expression of various mitochondrial enzymes ^19,34,35^ and HSPD1 ^20^ in NB cells. Here, we found that HSPD1 depletion in NB cells reduces MYCN expression, establishing a link between mitochondrial protein quality control and MYCN expression.

Previously, it was shown that pharmacological inhibition of mitochondrial translation induced the integrated stress response, leading to reduced MYCN expression due to reduced *MYCN* translation and sustained degradation ^8^. In addition, HSPD1 depletion was found to cause aggregation of mitochondrial ribosomal proteins and inhibit mitochondrial translation in HEK293T cells ^33^, which is expected to reduce MYCN protein levels as well. Apart from regulation at the translational level, the mitochondrial unfolded protein response (mtUPR) transcription factor ATF5 was shown to protect MYCN from FBXW7-mediated proteasomal degradation by binding FBXW7 in NB cells ^19^ . We found that in NB cells, MYCN stability was unaffected by HSPD1 depletion, whereas HSPD1 depletion reduced *MYCN* transcript levels. While we did not rule out the possibility that reduced cytosolic translation and sustained degradation rates in HSPD1 KD cells may have led to reduced total MYCN protein levels, the reduced *MYCN*-encoding mRNA does support the notion that HSPD1 regulates MYCN at the mRNA level.

The 3’ UTR of *MYCN* mRNA was found to regulate *MYCN* mRNA stability by serving as a binding site for MDM2 ^36^, which stabilizes *MYCN*, and the *MYCN* negative regulator miR-34a ^37^. Our RNA-seq data show that *MDM2* and *miR-34a* expression are increased in HSPD1 KD cells. We do not know the reason for the reduction in *MYCN* mRNA levels, as MDM2 and miR-34a have opposing effects on *MYCN* mRNA. In addition, we do not know if *MYCN* mRNA is more degraded or less transcribed in HSPD1 KD cells. Interestingly, HSPD1 KD was found to downregulate c-MYC transcription in prostate cancer by promoting an ATP crisis, leading to reduced beta-catenin signaling ^17^. It is possible that a similar HSPD1-MYCN axis exists in NB cells. Nevertheless, we found that HSPD1 KD reduces mitochondrial translation in NB cells, consistent with findings in HEK293 and yeast cells ^33^. Inhibition of mitochondrial translation was shown to reduce MYCN levels ^8^. It is therefore plausible that HSPD1 depletion leads to aggregation of mitochondrial ribosomal proteins, reduced mitochondrial translation, and reduced MYCN expression.

Using a xenograft model, we found that HSPD1 KD NB KELLY cells adapted to or were selected for upregulated MYCN expression without fully restoring HSPD1 expression in vivo. It was previously shown that NB cells do not adapt to doxycycline-mediated inhibition of mitochondrial translation ^8^, and the mechanism of adaptation identified here remains unknown. The finding that tumors generated by HSPD1 KD cells were less proliferative and more apoptotic suggests that the cells did not fully adapt to HSPD1 KD and that HSPD1 pro-tumorigenic functions are, in part, MYCN independent. These data support the idea that inhibiting HSPD1 may be a therapeutic opportunity for treating *MYCN*-amplified NB cells and that such treatment might benefit from a combination with apoptosis-inducing compounds.

## Supporting information

sup data, methods, full scan blots

