## Supplementary material for "HSPD1 promotes neuroblastoma by augmenting MYCN expression": sup data, methods, full scan blots

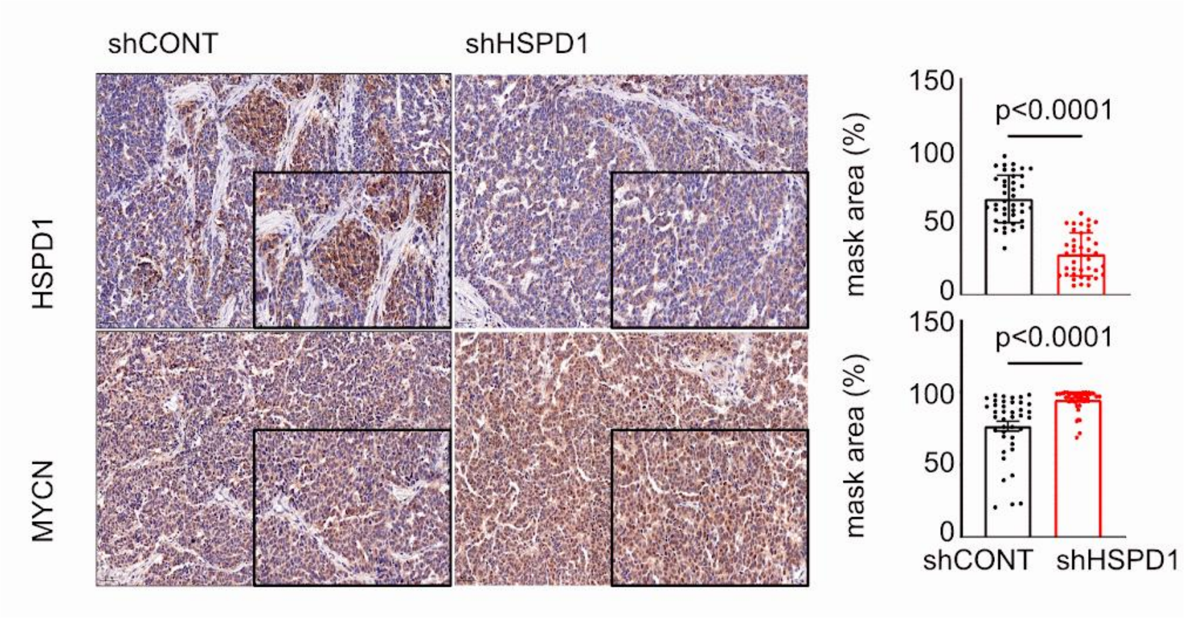


**Fig. S1. HSPD1 and MYCN levels in tumors.**
Tumors generated from HSPD1 KD and control KELLY cells were analyzed by IHC using anti-HSPD1 or MYCN antibodies. Staining was quantified as described in the Methods section and compared using Student's t-test.

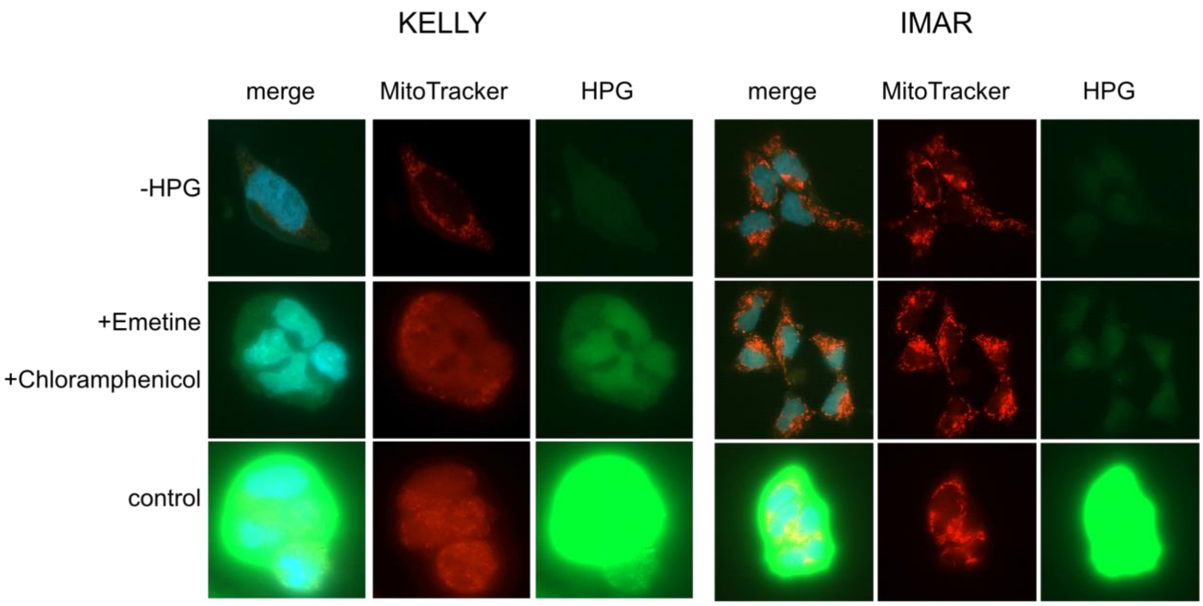


**Fig. S2. Controls for mitochondrial translation measurements.**
Experimental control images in shCtrl KELLY/ IMAR cells showing background fluorescence in the absence of label (-HPG), complete inhibition of both cytosolic and mitochondrial translation using combined emetine and chloramphenicol treatment (+Emetine +Chloramphenicol), and uninhibited total cellular translation (control). Scale bar =10 μm

### Blots - original scans

Fig. 2A

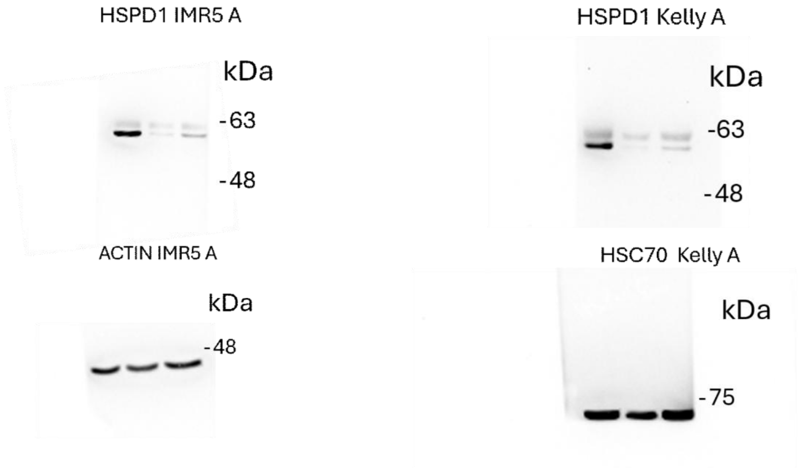


Fig. 5A

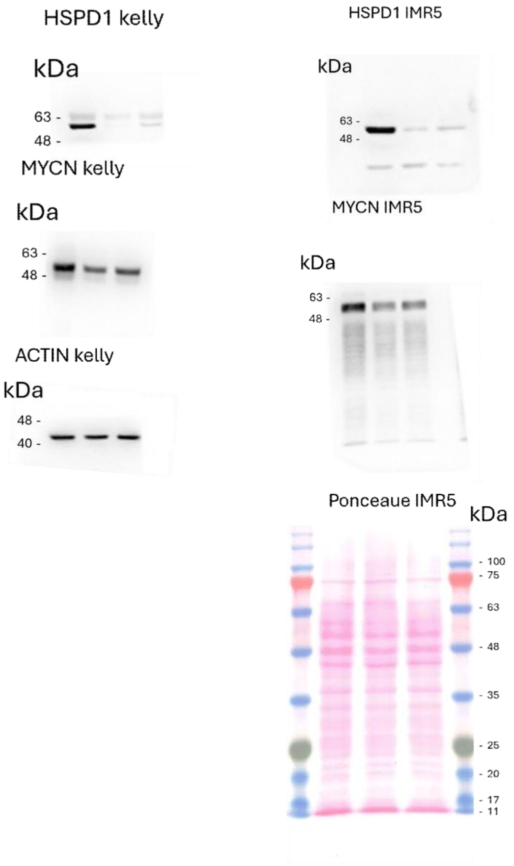

Fig. 5C

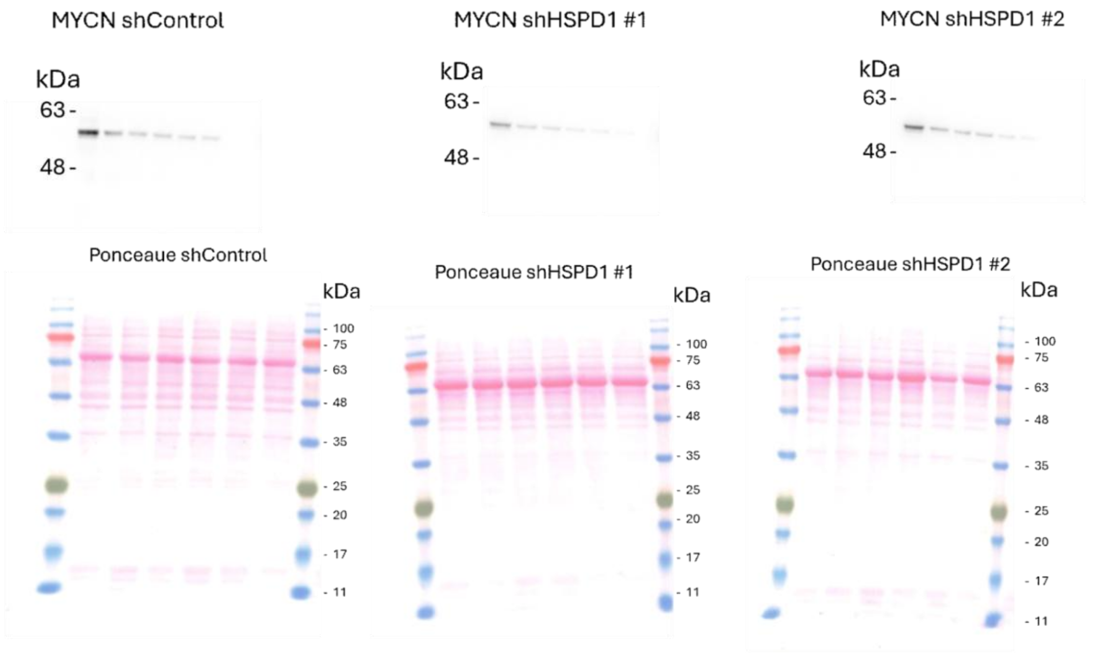


### Methods

#### Cell culture

KELLY cells were obtained from ATCC, and IMR-5 cells were a kind gift from Liron Grossman. KELLY and IMR-5 cells were cultured in Roswell Park Memorial Institute 1640 (RPMI-1640) medium. Media were supplemented with 10% fetal bovine serum (FBS), 1 mM sodium pyruvate, and antibiotic–antimycotic solution to create complete growth media. All cell lines were maintained under standard tissue culture conditions in a humidified incubator at 37 °C with 5% $CO_2$ and atmospheric oxygen.

#### Generation of stable cell lines for gene knockdown

To deplete HSPD1 we used CRISPR to generate heterozygotes and shRNA. Gene knockout of HSPD1 was performed using the SYNTHEGO Gene Knockout Kit v2. KELLY and IMR-5 cells were seeded into 6-well plates at a density of 200,000 cells per well and allowed to grow overnight. The next day, cells were transiently transfected with HSPD1-specific CRISPR guide RNAs pool (gRNAs) and CAS9 to a final concentration of 3µM, using Lipofectamine Cas9 Plus reagent and following the recommended protocol. HSPD1 protein levels were measured using western blot three days post-transfection, and three "pool-CRISPR-knockdown" populations were analyzed. WT cells were transfected similarly, without CAS9. Next, a single clone isolation protocol was performed to select clones with HSPD1 downregulation. Pool-CRISPR cells were seeded in 96 well plates at a concentration of 0.5 cells well. Filtered DMEM from proliferating WT cells was used in addition to fresh DMEM until small colonies were observed by light microscopy. Once clones filled a well, they were transferred to a larger well plate. Immunoblotting was used to validate the HSPD1 knockdown.

##### Lentiviral silencing

Two different CRISPR clones were chosen for HSPD1 knockdown using two independent (Short hairpin RNA) shRNA (SIGMA ALDRICH, MISSION shRNA, TRCN0000029446 and TRCN0000343952 for sh_1 and sh_2, respectively). For lenti virus generation, HEK293T Cells were grown in complete DMEM and transfected using CalFectin and a plasmid ratio of 1:2:3 (PAX2; pMD2.G; pLKO transfer vector). The media was changed after 24 hours, and the viruses were collected after 48 hours. The collected viruses were stored at -80°C. Prior to virus infection, cells were grown to 60% confluence on 6-well plates and then infected with the virus at a ratio of 1:10 (shCONT, sh#1, sh#2) and left for 24 hours in a 37°C incubator. Following infection, the containing medium was removed, replaced with a fresh medium, and incubated at 37°C. Puromycin (1 µg/ml) was used for selecting infected cells. Gene knockdown was validated using qRT-PCR and Western blot.

#### Quantitative Reverse Transcription Polymerase Chain Reaction (qRT-PCR)

Total RNA was extracted from cells using an RNA purification kit (Invitrogen), following the manufacturer's instructions. RNA concentration and quality were determined using a NanoDrop spectrophotometer. cDNA synthesis was performed using the cDNA synthesis kit (BioLabs), according to the manufacturer’s protocol, with approximately 800 ng of RNA as the input. For qPCR, reactions were carried out in a final volume of 12 µL, containing 6 µL of 2× SYBR Green Master Mix (PCR Biosystems), 1 µL of cDNA, 0.5 µM of forward and reverse primers (purchased from IDT), and 4 µL of nuclease-free water. PCR reactions were performed on iCycler Real-Time PCR System (Bio-Rad Laboratories) with the following cycling conditions: initial denaturation at 95°C for 10 minutes, followed by 46 cycles of 95°C for 10 seconds, 58°C for 30 seconds, and 72°C for 3 seconds. Finally, an elongation step was performed at 40°C for 10 minutes to ensure complete extension of any remaining single-stranded DNA. qRT-PCR data were acquired using the iCycler software, with threshold cycle (Ct) values determined from 4 replicate reactions. Relative gene expression was calculated using the $2^{(-\Delta\Delta Ct)}$ method, normalized to the expression of L32 as a housekeeping gene.

##### Western blot

Cells were cultured in a 10 cm dish to 90%-95% confluency and were lysed with RIPA lysis buffer supplemented with protease inhibitor (1:10000) and phosphatase inhibitor (1:10000). The lysates were sonicated for homogenization, and Pierce$^{TM}$ BCA Protein Assay Kit was used for protein quantification. A minimum of 10 µg of the protein lysate was subjected to sodium dodecyl sulfate-polyacrylamide gel electrophoresis (SDS- PAGE) and transferred onto nitrocellulose (BioTrace$^{TM}$, #66485) membranes. Membranes were stained with ponceau S and filmed before blocking with 5% skim milk powder in Tris buffer saline (TBS) for 1 hour and incubated with primary antibody diluted in blocking buffer overnight at 4°C. Next, the membranes were washed with TBS and incubated with HRP-linked secondary antibodies. Signals were determined using a chemiluminescence detection kit (Advansta, #K-12043-D20). ImageJ software was used to quantify band intensities.

#### Immunofluorescence and mitochondria staining

Cells were cultured in chambers. The next day, they were rinsed with phosphate-buffered saline (PBS) and fixed with 4% paraformaldehyde for 10 minutes at room temperature, followed by two additional washes. Cells were permeabilized with 0.1% tri-sodium citrate and 0.1% Triton X-100 in distilled water, pH 6, for 5 minutes and rinsed again. After 60 minutes of blocking with 0.5% bovine serum albumin (BSA), 5% goat serum, and 0.1% Tween-20 in PBS, cells were incubated overnight at 4°C with primary antibody diluted 1:200 or 1:1000 in blocking buffer. The next day, cells were washed three times with wash buffer (0.25% BSA, 0.1% Tween-20 in PBS), incubated for 1 hour with secondary antibody diluted 1:200 in blocking buffer at room temperature, then rinsed three more times. Cells were stained with DAPI for three minutes at room temperature, rinsed twice with PBS, and imaged. Nuclear mean intensity was measured and analyzed using ImageJ.

##### Mitochondrial Translation Assay

To evaluate mitochondrial translation, newly synthesized mitochondrial proteins were quantified using L-homopropargylglycine (HPG; Click-iT HPG Alexa Fluor 488 Protein Synthesis Assay Kit, Thermo Fisher Scientific, Cat. No. C10428). IMR/ Kelly NB cell lines shControl / HSPD1 knockdown (KD) were seeded in 8-well chamber slides 24 h prior to treatment.

To deplete endogenous methionine, culture media were replaced with methionine-free starvation medium for a 15 min pre-incubation period at 37 °C, with specific translation inhibitors added during this pre-incubation: emetine (50 µg/mL) to selectively inhibit cytosolic translation, or emetine (50 µg/mL) and chloramphenicol (100 µg/mL) to block both cytosolic and mitochondrial translation. Following the 15 min pre-incubation, HPG was added directly into the medium while maintaining the presence of the respective inhibitors. Baseline background fluorescence was obtained by incubation in methionine-free starvation medium without HPG addition.

After 30 min of HPG labeling, MitoTracker™ Red CMXRos (Thermo Fisher Scientific, Cat. No. M46752) was added directly into the medium to a final concentration of 200 nM and incubated for an additional 30 min, resulting in a total HPG labeling period of 1 h. Following incubation, cells were fixed with 3.7% formaldehyde in PBS for 15 min at room temperature, permeabilized with 0.5% Triton® X-100 in PBS, and reacted with Alexa Fluor 488 azide according to the manufacturer’s instructions.

##### Imaging and Mitochondrial Signal Quantification

Fluorescence images were captured using a Zeiss *Celldiscoverer* 7 automated microscope. Image processing, segmentation, and quantitative analysis were conducted using *CellProfiler* software (Version 4.x).

To correct for uneven illumination and local background, raw fluorescence images from both the MitoTracker™ Red CMXRos and Alexa Fluor 488 (HPG) channels were processed using the *Correct Illumination Calculate* and *Correct Illumination Apply* modules. Mitochondrial networks were segmented from the background-corrected MitoTracker™ Red channel using the *Identify Primary Objects* module, applying an adaptive two-class Otsu thresholding method (threshold correction factor of 1.0) with a pixel diameter range optimized to exclude non-specific staining artifacts. A binary mitochondrial mask was generated using the *Mask Objects* module. The *Measure Object Intensity* module was then applied to quantify the mean fluorescence intensity of the Alexa Fluor 488 (HPG) signal strictly within the boundaries of the mitochondrial mask. Data were exported using the *Export To Spreadsheet* module. Signal intensities were normalized relative to the untreated control condition (ShControl with emetine), which was set to a maximum reference value of 1.

**Total Translation Assay**

To evaluate total cellular translation, newly synthesized proteins were quantified using L-homopropargylglycine (HPG; Click-iT HPG Alexa Fluor 488 Protein Synthesis Assay Kit, Thermo Fisher Scientific, Cat. No. C10428). IMR NB cell lines shControl / HSPD1 KD were seeded in 8-well chamber slides 24 h prior to treatment.

To deplete endogenous methionine, culture media were replaced with methionine-free starvation medium for a 15 min pre-incubation period at 37 °C, either in the absence of inhibitors (total translation) or in the presence of combined translation inhibitors: emetine (50 µg/mL) and chloramphenicol (100 µg/mL) to block overall cellular translation. Following the 15 min pre-incubation, HPG was added directly into the medium while maintaining the presence or absence of the inhibitors, resulting in a total HPG labeling period of 1 h. Baseline background fluorescence was obtained by incubation in methionine-free starvation medium without HPG addition. Following incubation, cells were fixed with 3.7% formaldehyde in PBS for 15 min at room temperature, permeabilized with 0.5% Triton® X-100 in PBS, and reacted with Alexa Fluor 488 azide according to the manufacturer's instructions.

**Imaging and Total Signal Quantification**

Fluorescence images were captured using a Zeiss *Celldiscoverer* 7 automated microscope. Image processing, segmentation, and quantitative analysis were conducted using *CellProfiler* software (Version 4.x).

To correct for uneven illumination and local background, raw fluorescence images from the Alexa Fluor 488 (HPG) channel were processed using the *Correct Illumination Calculate* and *Correct Illumination Apply* modules. Cellular boundaries were segmented from background-corrected images using the *Identify Primary Objects* module, applying an adaptive two-class Otsu thresholding method (threshold correction factor of 1.0) to define cellular regions. The *Measure Object Intensity* module was then applied to quantify the mean fluorescence intensity of the Alexa Fluor 488 (HPG) signal within the segmented cells. To account for differences in cell density across fields of view, total signal intensity was normalized to the number of nuclei per image. Data were exported using the *Export To Spreadsheet module*.

**Statistical Analysis**

Statistical analyses were performed using GraphPad Prism software (version 11.0, GraphPad Software, San Diego, CA, USA). Data were evaluated using a one-way Analysis of Variance (ANOVA) followed by Dunnett's multiple comparisons test to compare experimental conditions against the control group. Statistical significance was set at $p < 0.05$.

#### Cycloheximide Chase Analysis

Cells were seeded one or two days prior to the experiment, followed by incubation with a final concentration of 50µg/ml cycloheximide (CHX) in RPMI-1640 to inhibit protein synthesis. Subsequently, cells were harvested at different time points after CHX treatment (0, 1, 1.5, 2, 2.5, 3 hours) for protein extraction, which was analyzed by Western blot. The MYCN bands were quantified using ImageJ software, with actin bands or Ponceau staining used for normalization. The MYCN half-life was determined by transforming the data to the natural logarithm (ln), performing linear regression for each cell line, and dividing ln (0.5) by the slope of each regression line.

#### Mouse Xenograft Model

All animal procedures complied with the institutional animal care and use committee and relevant guidelines at Ben-Gurion University (permission number: 71-10-2023-E). Mice were housed in specific-pathogen-free (SPF) conditions at the Ben-Gurion University facility. All efforts were made to minimize animal suffering and ensure humane treatment.

A mouse xenograft model was employed to investigate the effects of HSPD1 knockdown on tumor growth. A total of $8 \times 10^6$ KELLY control cells or HSPD1 knockdown cells, resuspended in 100 µL PBS, were injected subcutaneously into the left and right flanks of each 8-week-old female NOD.CB17-Prkdc$^{scid}$ mouse ($n$ = 5 mice per group). Tumor development was monitored over 37 days, with caliper measurements taken at regular intervals. At the end of the experiment, mice were euthanized using $CO_2$ asphyxiation as IACUC guidelines. Tumors were excised and processed for downstream analysis. Tissue was fixed in 4% paraformaldehyde (PFA) and processed for IHC.

All mouse work was performed in accordance with the institutional animal care use committee and relevant guidelines at the Ben-Gurion University, with protocol IL80-09-2023E.

#### IHC

human samples

Formalin-fixed, paraffin-embedded TMA sections were analyzed for MYCN and HSP60 expression. In brief, tissue sections were incubated in Tris EDTA buffer (cell conditioning 1; CC1 standard) at 95 °C for 1 hour to retrieve antigenicity, followed by incubation with MYCN (ab198912, 1:150) or HSP60 antibody (Sigma PLA0269, 1:500) for 1 hour. Slides were then incubated with secondary antibody (Jackson Laboratories) with 1:500 dilution, followed by Ultramap HRP and Chromomap DAB detection. Intensity scoring was performed on a common four-point scale. Descriptively, 0 represents no staining, 1 represents low but detectable degree of staining, 2 represents clearly positive staining, and 3 represents strong expression. Expression was quantified as H-Score, the product of staining intensity, and % of stained cells.

The TMAs including 33 NB used in the study were obtained from the Children's Oncology Group (COG). H&E stained slides from these tissue blocks were annotated for desired areas and the slides were then used as guides for core selection. TMAs were constructed on a Beecher Manual Tissue Arrayer I (MTA I). For all TMAs, 1mm cores of normal FFPE tissue were punched from a donor block and arrayed into a recipient paraffin block. Cores were placed into the recipient block following a coordinate grid map. Each tissue or tumor type was represented by three cores when possible.

Mouse samples

Tissues were fixed in 4% paraformaldehyde (PFA) at room temperature for up to 24 hours, dehydrated, and embedded in paraffin. Tissue sections were deparaffinized using xylene, and endogenous peroxidase activity was blocked by treating with 3% hydrogen peroxide ($H_2O_2$) for 20 minutes. Subsequently, sections were rinsed with water for 5 min. Antigen retrieval was carried out in citrate buffer (pH 6) at 99.9 °C for 15 minutes. The sections were then blocked for 1 hour at room temperature with a blocking solution composed of phosphate-buffered saline (PBS), 0.1% Tween, and 5% bovine serum albumin (BSA). Following blocking, the sections were incubated overnight at 4 °C with the primary antibody Ki67 (1:200; ab16667), Cleaved caspase 3 (1:200; CST-9664S), HSP-60 (1:100), and N-MYC (1:100) diluted in blocking solution. On the next day, the slides were washed three times with PBS-Tween (PBS-T), and color detection was performed using an ABC kit (VECTASTAIN, Cat. VE-PK-6200) as per the manufacturer's instructions. Sections were counterstained with hematoxylin and mounted using Micromount mounting medium (Leica, Cat. 380-1730). All slides were digitized using a Panoramic Scanner (3DHISTECH, Budapest, Hungary), and image analysis was performed with Qupath-0.2.3 software. Tumor tissue fields for analysis were selected by a blinded investigator. Cells in the selected fields were detected using Qupath-0.2.3 software and classified as positive or negative for DAB staining based on thresholds determined independently by two blinded investigators. Cell detection criteria and threshold values were standardized across all slides. To confirm the specificity of the staining and analysis thresholds, a matched negative control was used. This control was processed identically but without incubation with the primary antibody, ensuring all secondary antibody development procedures were followed.Immunohistochemistry (IHC) and H-scores

Formalin-fixed, paraffin-embedded TMA sections were analyzed for MYCN and HSP60 expression. In brief, tissue sections were incubated in Tris EDTA buffer (cell conditioning 1; CC1 standard) at 95C for 1 hour to retrieve antigenicity, followed by incubation with MYCN (ab198912, 1:150ss) or HSP60 antibody (Sigma PLA0269, 1:500) for 1 hour. Slides were then incubated with secondary antibody (Jackson Laboratories) with 1:500 dilution, followed by Ultramap HRP and Chromomap DAB detection. Intensity scoring was performed on a common four-point scale. Descriptively, 0 represents no staining, 1 represents low but detectable degree of staining, 2 represents clearly positive staining, and 3 represents strong expression. Expression was quantified as H-Score, the product of staining intensity, and % of stained cells.

##### Crystal violet staining

Cell viability was evaluated using crystal violet staining. The cells were seeded in 12-well plates at a density of 10,000 cells per well and cultured for five days under standard conditions. After incubation, cells were gently rinsed with phosphate-buffered saline (PBS) and allowed to air-dry for 5 minutes. The cells were then stained with crystal violet solution for 30 minutes at room temperature. Excess stain was rinsed off with distilled water, and the plates were allowed to dry completely. The dye was then solubilized with 10% acetic acid, and absorbance was measured at 595 nm using a microplate reader to quantify relative cell viability.

#### RNA sequencing

Total RNA was purified from the TRIzol fractions collected during the ribosome profiling. To separate the phases, chloroform was added to the samples and centrifuged at 20,000 × g for 15 minutes at 4 °C. The aqueous phase was carefully transferred to a new tube and adjusted with 3 M sodium acetate to a final concentration of 0.3 M, followed by the addition of 2 µL GlycoBlue and an equal volume of isopropanol. The mixture was vortexed briefly and incubated at –20 °C overnight to precipitate RNA.The following day, RNA was pelleted by centrifugation at 20,000 × g for 45 minutes at 4 °C, washed once with 70% ethanol, air-dried, and resuspended in 10 mM Tris-HCl (pH 7.0) . RNA integrity was determined on a QIAxcel device (QIAGEN, QIAxcel RNA QC Kit v2.0), and concentration was determined using a QuantiFluor® RNA System (Promega, #E3310) on a Qubit™ Flex Fluorometer (Invitrogen). mRNA enrichment was conducted from 0.5µg of total RNA using the NEBNext® Poly(A) mRNA Magnetic Isolation Module (New England Biolabs, #E7490). Stranded RNA-seq libraries were constructed using NEBNext® Ultra™ II Directional RNA Library Prep Kit for Illumina® (New England Biolabs, #E7760). The molarity of libraries was determined using QIAxcel (QIAxcel DNA High Sensitivity Kit) and QuantiFluor® dsDNA System (Promega, # E2670) on a Qubit™ Flex Fluorometer (Invitrogen).  Libraries were sequenced (150PE) on a NovaSeq X (Illumina).

**Table 1. List of materials used in this study**

| Material | Source | Identifier |
|---|---|---|
| RPMI 1640 Medium | Biological Industries | 01-100-1A |
| DMEM, high glucose | Gibco | Cat#11965092 |
| Fetal bovine serum (FBS) | ThermoFisher Scientific | Cat#10270106 |
| sodium pyruvate solution | Biological Industries | Cat#03-042-1 |
| antibiotic–antimycotic 100X | TOKU-E | Cat#A045 |
| Gene Knockout Kit v2 | SYNTHEGO | Cat#111 |
| Lipofectamine CRISPRMAX Cas9 transfection reagents | ThermoFisher Scientific | Cat#CMAX00001 |
| Opti-MEM I reduced serum medium. | ThermoFisher Scientific | Cat#31985062 |
| DAPI Fluoromount-G® | Southern Biotech | Cat#0100-20 |
| Dimethyl sulfoxide (DMSO) | Sigma-Aldrich | Cat#D8418 |

| | | |
|---|---|---|
| Ponceau S solution | Sigma-Aldrich | Cat#P7170-1L |
| CalFectin™Mammalian Cell Transfection Reagent | SignaGen Laboratories | Cat#SL100478 |
| Protease Inhibitor Cocktail | Sigma-Aldrich | Cat#P8340 |
| PhosSTOP™ | Roche | Cat#4906837001 |
| Crystal violet | Sigma-Aldrich | Cat#C0775 |
| Cycloheximide (CHX) | Sigma-Aldrich | Cat#01810 |
| PBS | Invitrogen | Cat# 10010-023 |
| Triton X-100 | Sigma-Aldrich | Cat# 000010 |

**Table 2. Antibodies used in this study**

| Antibody | Source | Identifier |
|---|---|---|
| MYCN (WB: 1:1000) (IF 1:2000) | Santa Cruz Biotechnology | Cat# sc-53993 |
| MYCN (IHC human samples; 1:150) | Abcam | Cat# ab198912 |
| HSPD1 (WB: 1:1000) | Santa Cruz | Cat#sc-13115 |
| HSPD1 (IF: 1:1000) | Cell Signaling Technology | Cat#12165 |
| HSPD1 (IHC 1:100) | Sigma | Cat# PLA0269 |
| Phospho-S6 (WB: 1:1000) | Cell Signaling Technology | Cat# 2215S |

| Antibody | Source | Identifier |
| --- | --- | --- |
| MYCN (WB: 1:1000) (IF 1:2000) | Santa Cruz Biotechnology | Cat# sc-53993 |
| MYCN (IHC human samples; 1:150) | Abcam | Cat# ab198912 |
| Anti-mouse IgG (WB: 1:1500) | Cell Signaling Technology | Cat#7076 |
| Anti-rabbit IgG (WB: 1:1500) | Cell Signaling Technology | Cat#7074 |
| Actin (WB: 1:10000) | Merck | Cat# MAB1501 |
| HSC-70 (WB: 1:5000) | Santa Cruz | Cat#sc-7298 |
| Alexa 488 anti-Rabbit IgG (IF 1:200) | ENCO | Cat#111-545-144 |
| Alexa 594 anti-Mouse IgG (IF 1:200) | ENCO | Cat#115-585-062 |
| Ki67 | Abcam | Cat# ab16667 |
| Cleaved caspase 3 | Cell Signaling Technology | Cat# CST-9664S |

**Table 3. shRNA sequences that were used**

| | |
| --- | --- |
| shControl | CCTAAGGTTAAGTCGCCCTCGCTCGAGCGAGGGCGACTTAACCTTAGG |
| shHSPD1 #1 | CCGGCCTGCTCTTGAAATTGCCAATCTCGAGATTGGCAATTTCAAGAGCAGGTTTTT |
| shHSPD1 #2 | CCGGGCAATGACCATTGCTAAGAATCTCGAGATTCTTAGCAATGGTCATTGCTTTTT |

**Table 4. Plasmids used in this study**

| Plasmid | Source | Identifier |
| --- | --- | --- |

| | | |
|---|---|---|
| pMD2.G | Addgene | Cat#12259 |
| psPAX2 | Addgene | Cat#12260 |

**Table 5. Buffers used in this study**

| Buffer | Component and final concentrations |
|---|---|
| RIPA lysis buffer | 150 mM NaCl; 50 mM Tris pH = 8.0; 1**%** Triton X-100; 0.5% sodium deoxycholate; 0.1% SDS |
| TBS (WB) | 20 mM Tris HCl, pH 7.4, 150 mM NaCl |
| Running buffer (WB) | 25 mM Tris base, 192 mM Glycine, 0.1% SDS, pH=8.3 |
| Transfer Buffer (WB) | 12 mM Tris base, 196mM Glycine, pH=8.3 |
| Permeabilization Buffer (IF) | 0.1% tri-sodium citrate, 0.1% Triton X-100 in distilled water, pH 6.0 |
| Blocking Buffer (IF) | 0.5% bovine serum albumin (BSA), 5% goat serum, 0.1% Tween-20 in PBS |
| Wash Buffer (IF) | 0.25% BSA, 0.1% Tween-20 in PBS |
| 5X Sample buffer | 250 mM Tris, 40% Glycerol, 10% SDS, 0.25% bromophenol blue, pH 6.8 |

**Table 6. Experimental models Organism/Strain**

| Organism/Strain | Source | Identifier |
|---|---|---|
| Mouse: NOD *SCID* Gamma *Prkdc*$^{scid}$ | The Jackson Laboratory | RRID: IMSR_JAX:001303 |
